# A Priming Circuit Controls the Olfactory Response and Memory in *Drosophila*

**DOI:** 10.1101/2025.03.06.641741

**Authors:** He Yang, Yang Jiang, Samuel Kunes

**Affiliations:** Department of Molecular and Cellular Biology, Harvard University, Cambridge, Massachusetts, USA; Program in Neuroscience, Department of Neurobiology, Harvard Medical School, Boston, Massachusetts, USA; Janelia Research Campus, Howard Hughes Medical Institute, Ashburn, Virginia, USA

## Abstract

Priming is a process by which exposure to a stimulus affects the response to a subsequent stimulus in humans. In this study we found that in *Drosophila*, a prior encounter with an aversive stimulus results in enhanced preference for a following novel odor, while an appetitive stimulus leads to reduced preference for a new odor. This priming behavior of flies relies on the well-studied olfactory memory circuits including Kenyon cells (KC), dopaminergic neurons (DANs), and mushroom body output neurons (MBONs). Aversive stimulus results in increased odor responses in reward DANs that innervate the γ4 lobe of the mushroom body (MB) and decreased odor responses in a γ4γ5-innvervating repulsive MBON. We concluded that these neurons are required for the priming effects in flies. These results characterized the newly found priming behavior in flies and demonstrated the sheer influence of unconditioned stimulus on odor perception during associative learning.

## Introduction

Animals receive rich sensory stimuli in a changing environment. To avoid danger and search for food, they must discriminate different sensory cues, memorize the associated outcomes, and update their behaviors based on the past experiences. Olfactory learning has been extensively studied in fruit fly *Drosophila melanogaster*, which can be conditioned to associate an odorant as a conditioned stimulus (CS) with an aversive or appetitive unconditioned stimulus (US) including electric shocks (Quinn et al., 1974; Tully and Quinn, 1985) and sugar feeding (Tempel et al., 1983). Various forms of conditioning have been observed in *Drosophila* by modifying the temporal relationship between CS and US during the training paradigm, including trace conditioning (Galili et al., 2011) and backward conditioning (Tanimoto et al., 2004). However, the sheer impact of US itself on sensory perception has not been well studied. Here we find that exposure to an aversive or appetitive US alone results in enhanced or decreased preference, respectively, for a subsequently presented novel odor in *Drosophila*. We characterize this modulation of preference as a priming effect, a phenomenon previously defined as the influence of one stimulus on the perception of or response to a subsequent stimulus, typically occurring without conscious awareness or deliberate processing (Murphy and Zajonc, 1993). While priming effects have been investigated in human subjects within social science research (Bargh, 2006), they are infrequently reported in non-human animals (Brodbeck, 1997; McMahon and Olson, 2007; Amitai et al., 2013; Sivroni et al., 2023), and typically require prior associative training protocols to account for the animals’ lack of verbal ability.

While the molecular or cellular electrophysiology underlying priming behavior has not been studied in any animal, it was shown that priming effects require memory formation (Beer and Diehl, 2001). To map the priming circuits in flies, we focused on neurons underlying olfactory learning and memory. In the MB, the center of learning and memory for insects (Heisenberg et al., 1985; De and Heisenberg, 1994; McGuire et al., 2001; Menzel, 2014), specific odor identities are encoded by MB intrinsic KCs sparsely (Wang et al., 2004; Turner et al., 2008; Campbell et al., 2013; Caron et al., 2013), and then converged onto MBONs, which represent more abstract information including the valence of an odor based on learning experiences (Owald et al., 2015; Hige et al., 2015a; Yamazaki et al., 2018). MBONs eventually activate downstream motor circuits to guide the avoidance or approach behaviors in flies. In each hemisphere, KCs are coarsely grouped into α/β, α’/β’, and γ classes based on birth order and adult axonal projection patterns (Crittenden et al., 1998; Lee et al., 1999). The MB lobes can be further segmented into 15 anatomically discrete compartments, each innervated by different subsets of MBONs and DANs. Together, these neurons and their connections provide the anatomical architecture for the formation and retrieval of different memory types (Tanaka et al., 2008; Mao and Davis, 2009; Aso et al., 2014a; Cohn et al., 2015).

The plasticity of the synapses between KCs and MBONs is modulated presynaptically by DANs that innervate their axons into the corresponding compartments of MB lobes. DANs are classified into 21 cell types (Aso et al., 2014a; Li et al., 2020), some of which are activated by US and thus convey the punishment or reward signals to animals. In general, DANs of the paired posterior lateral 1 (PPL1) cluster are punishment reinforcing, whereas protocerebral anterior medial (PAM) DANs usually encode reward (Schwaerzel et al., 2003; Claridge-Chang et al., 2009; Aso et al., 2010; Aso et al., 2012; Kirkhart and Scott, 2015; Jacob and Waddell, 2020). Different subsets of DANs innervate distinct, non-overlapping compartments of MB lobes in a robust pattern across individual flies, feeding the US information onto KC fibers which encode the odor identities previously neutral to flies, and thus updating the memories (Aso et al., 2010; Liu et al., 2012; Aso and Rubin, 2016). Outside the MB, some MBON axons also project to connect with DAN dendrites to form a recurrent loop (Eschbach et al., 2020; Li et al., 2020; Scheffer et al., 2020), together coordinating the plasticity of MBONs, DANs, and KCs across multiple MB compartments, giving rise to diversified olfactory responses and memory dynamics in fruit flies.

In this study, we investigated the bidirectional priming effect of aversive and appetitive US on the following odor responses in flies and found plasticity of odor responses in subsets of DANs and MBONs which innervate the γ4-MB. We also demonstrated that these neurons are required for the priming behavior. For the first time, we observed the impact of US alone on olfactory valence in flies. Our study broadened the understanding of olfactory associative learning by highlighting a previously unexplored component of the short-term memory in *Drosophila*.

## Results

### Preference for novel odors in *Drosophila* is altered by electric shocks

Over decades, olfactory memory in *Drosophila* has been tested and quantified by a canonical Pavlovian conditioning paradigm using a T-maze (Tully and Quinn, 1985). A group of flies first receive foot shocks from copper electrodes in a tube filled with a CS+ odor, then rest in the tube without shocks with the presence of a CS-odor. They are later transferred into a T-maze at the midpoint to choose between the two previously experienced odors which filled each arm. The performance index (PI) quantifies the learning and memory score of those flies, calculated as the fraction of flies that approach CS-subtracted by that approach CS+. This aversive olfactory memory was previously considered to result only from the flies’ repulsion to CS+ after shock pairing. However, recent studies (Jacob and Waddell, 2020) and our experiments have allowed us to focus on the CS-memory component by introducing a third odor into the test phase (Figure 1—figure supplement 1A-B). We found that after shock training, wildtype 2U flies showed significantly increased preference for CS-compared to either a third, novel odor or pure air (Figure 1—figure supplement 1C-F). The aversive olfactory memory could be considered as bipartite, consisting of a repulsion memory to CS+ and an attraction memory to CS-(Figure 1—figure supplement 1G-K). These results suggest that some neural circuits underlying appetitive memory are also likely to be involved in the formation and maintenance of the plasticity after aversive conditioning.

During shock training, CS-was delivered to flies but not directly paired with any US simultaneously. To test the necessity of the presence of CS-during training for forming attraction memory, we further modified the conditioning paradigm by removing CS-from training (Figure 1A). Wildtype flies shocked with CS+ pairing alone (1x 60s) were tested 1min later to choose between pure air and a novel chemical odor. Like in canonical conditioning, the odor paired with shock (Odor A) and the one used for test (Odor B) were swapped between 3-octanol (OCT) and 4-methylcyclohexanol (MCH), with the PI scores averaged between a pair of reciprocal experiments to cancel the bias of the naïve odor preference. Flies raised in the same food vial were randomly separated into two groups, one trained in a tube with electric foot shocks, another mock trained in an identical tube without shock. ΔPI of each food vial was calculated as the difference in PI between shocked and unshocked groups to minimize the effects of innate odor preference. Naturally, the chemical odors used in these experiments are both repulsive to flies compared to air, but when comparing the preference between mock trained and shock experienced flies, we found that shocked flies demonstrated a significantly enhanced preference for the novel odor with a ΔPI of approximately 0.15, despite this odor had not been conditioned in any way during training. We refer to this non-conditioning behavior as priming, because here shock stimuli altered the flies’ reactions to a following, non-specified odor.

**Figure 1.**
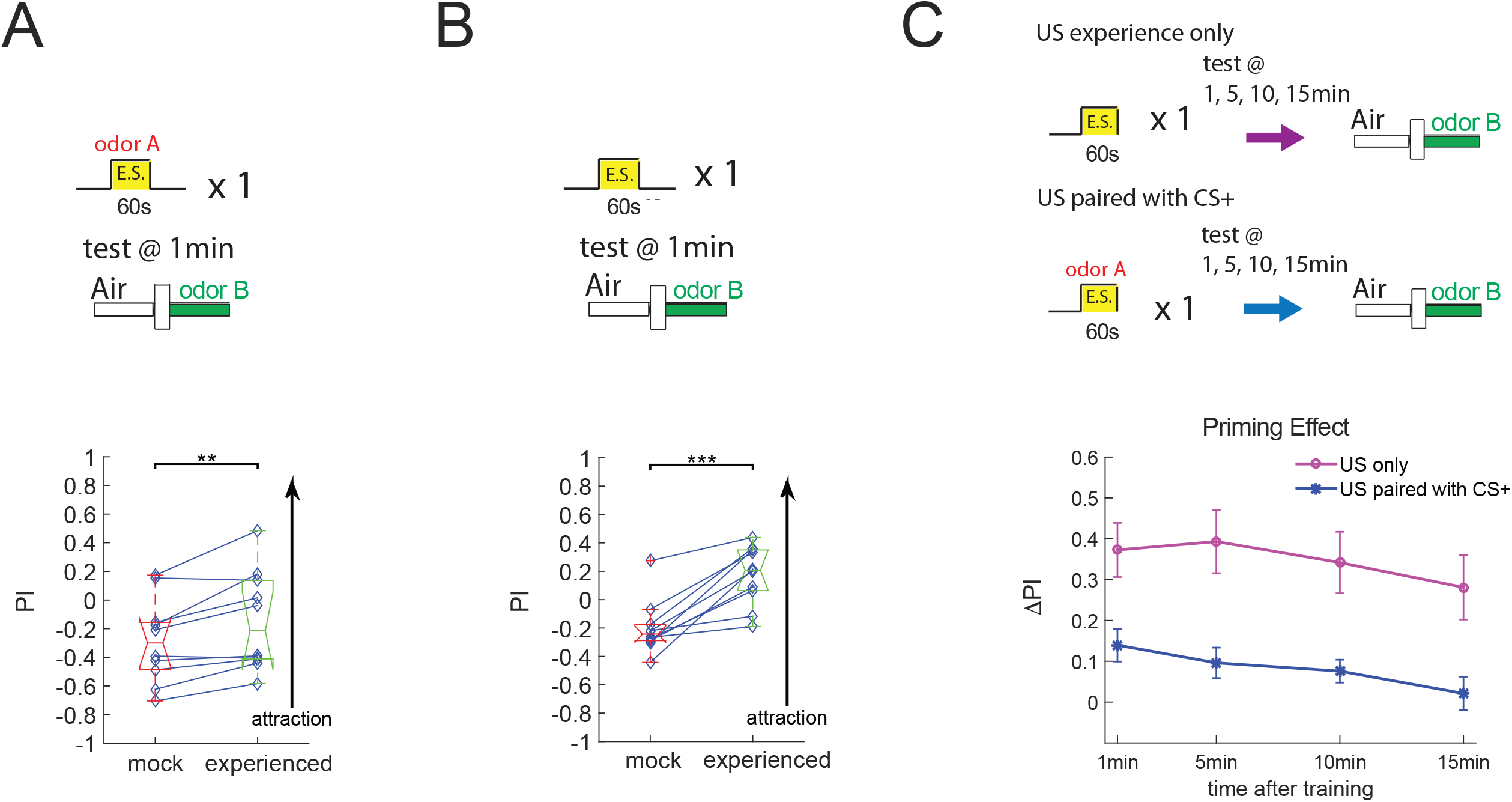
Shock priming changes flies’ preference for novel odors. (A-B) Electric shocks enhance the flies’ preference for a subsequent novel odor. In each panel, the training and test diagram is shown on the top, and the performance indices of mock-trained and primed flies are at the bottom. (A) Electric shocks paired with CS+ increased the preference for a novel odor when pure air was used as reference (n=10; paired t-test; ** *P* < 0.01). (B) Shocks alone also induced attraction to the novel odor (n=10; paired t-test; *** *P* < 0.001). (C) Enhanced preference for novel odors induced by shock experience lasts for minutes. Top: Schematic of priming with or without coincident odors. Bottom: Time course of the PI scores tested at different time points following training. ΔPI = primed PI – mock PI.

A more simplified priming paradigm was further explored with the removal of CS+ odor in addition to CS-(Figure 1B). In this setup, flies only received 1min of shock pulses before they were tested to choose between pure air and a novel odor. Results showed that shocked flies developed an even stronger preference for the tested odor compared to unshocked flies, with a ΔPI of approximately 0.4. We used this protocol as the major training method for most of the following experiments because the resultant preference change is more significant with a simpler training model. Flies in these assays were only shocked but not conditioned, aligning more closely with the definition of priming.

To understand how long the priming effect lasted, we trained different groups of flies by either electric shocks with CS+ pairing or shocks alone, and tested their relative preferences for the new odor at 1, 5, 10, or 15min after training (Figure 1C). We found that though diminished over time, this priming effect remained high 15min after 1min of shock pulses alone, while the effect with CS+ pairing appeared milder and lasted shorter. This relatively long-lasting impact of shock priming on odor valence rules out the interpretation of the priming effect as a backward conditioning between electric shocks and the odor presented later in the test, because flies cannot be conditioned with such a long temporal gap. The robust and long-lasting priming effect in flies implies profound changes in the underlying neural circuitry.

### Optogenetically substituted US induce both aversive and appetitive priming

Artificial activation or suppression of specific subsets of DANs by optogenetic or thermal approaches is sufficient to replace real punishment or reward to induce different forms of memories (Schroll et al., 2006; Claridge-Chang et al., 2009; Masek et al., 2015; Yamagata et al., 2015; Yamagata et al., 2016; Waddell, 2016). To determine whether US-substitution is sufficient to induce priming, we quantified the priming effect using an automated optogenetic olfactory circular arena in which flies could stay during both training and test, while different odors were delivered into specific quadrants of the arena as programmed (Aso et al., 2016). We first expressed the red-shifted channelrhodopsin, CsChrimson (Klapoetke et al., 2014), in punishment-encoding PPL1-γ1pedc DANs labeled by MB320C (Aso et al., 2016), and activated these neurons in approximately 20 flies by 1min of red LED pulses illuminating the entire arena with only air presented (Figure 2A). After another 1min of rest, the flies were tested to choose between pure air and odor OCT. We calculated the PI to quantify their preferences for OCT over air. As control, another group of flies raised from the same food vial underwent mock training without red LED. Compared to mock trained animals, flies that received light activation showed a significantly higher PI score, indicating an enhanced preference for OCT, which is consistent with the behaviors of the foot-shocked flies. As a genetic control, flies carrying UAS-CsChrimson and empty split-GAL4 (+/CsChrimson; Dionne et al., 2018), which had CsChrimson barely expressed in any neuron, showed no difference in their OCT preference between primed and mock-trained groups. We calculated a priming score to measure the priming effect, which was the ΔPI by subtracting the mock PI from the primed PI for each pair of experiments of flies from the same food vial. We concluded that after activating punishment DANs, these flies indeed showed a significantly higher priming score than the empty driver controls (Figure 2B).

**Figure 2.**
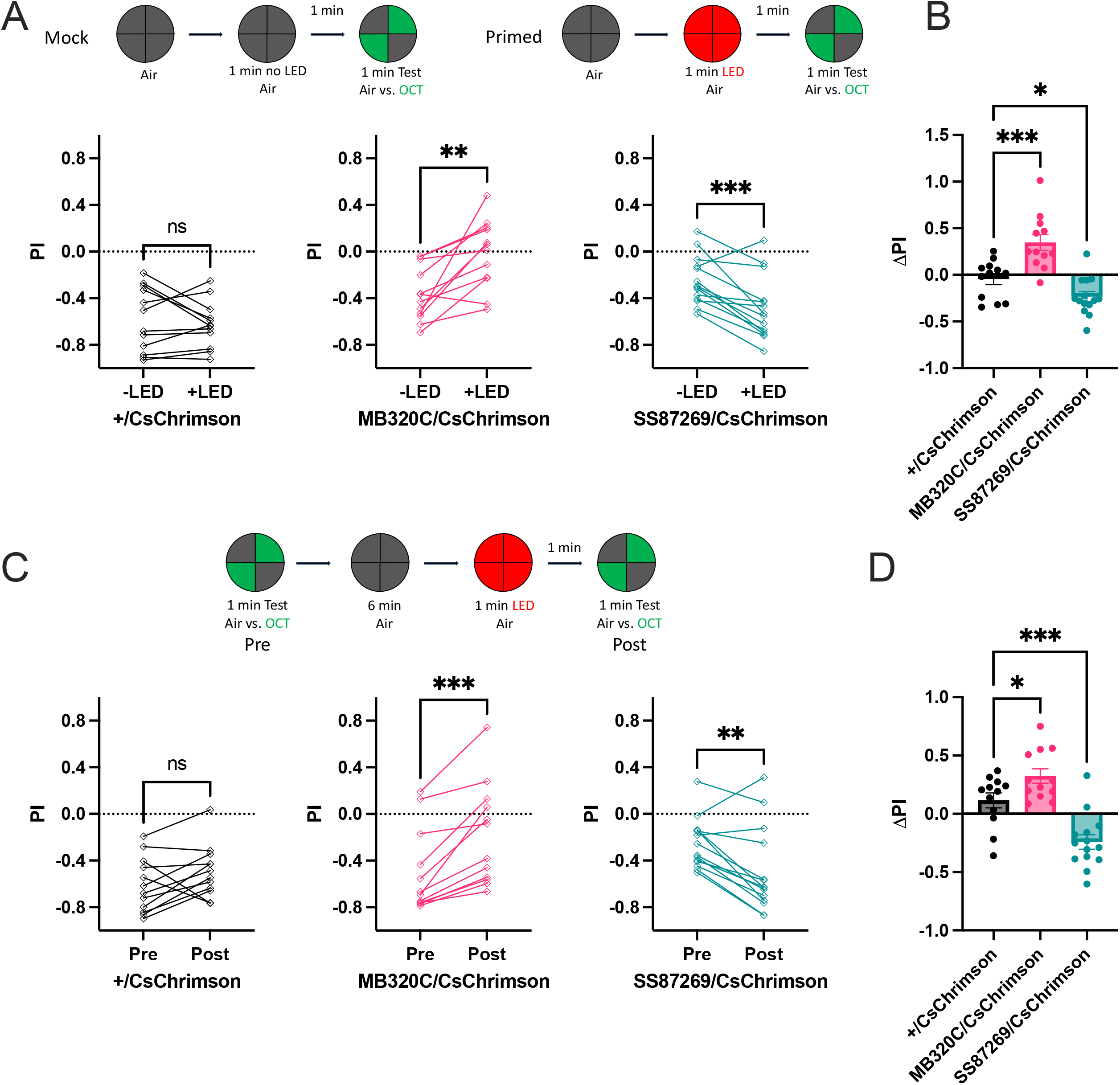
Both aversive and appetitive artificially substituted US induce priming effects. (A-B) Optogenetic substitution of US results in different odor preference compared to naïve flies. (A) Top: schematic of experimental protocol. Bottom: PI scores of mock-trained (-LED) flies comparing with primed (+LED) ones. From left to right: optogenetic activation of UAS-CsChrimson expressed in empty split-GAL4 (+, control, n=12), PPL1-γ1pedc DANs (MB320C, aversive cue, n=12), and sugar sensory neurons (SS87269, appetitive cue, n=14). (B) Activation of aversive or appetitive neurons induces priming bidirectionally. ΔPI = PI_+LED_ – PI_-LED_. (C-D) Flies change odor preference after optogenetic activation. (C) Top: experimental protocol. Bottom: PI scores of post-activation tests compared with pre-activation tests. From left to right: UAS-CsChrimson expressed in empty split-GAL4 (+, n=12), PPL1-γ1pedc DANs (MB320C, n=12), and sugar sensory neurons (SS87269, n=14). (D) Activation of aversive or appetitive neurons induces bidirectional priming. ΔPI = post PI – pre PI. (A and C: paired t-test within each genotype. B and D: one-way ANOVA. All ΔPI shown as mean ± SEM. The number of asterisks indicates the significance level: * *P* < 0.05; ** *P* < 0.01; *** *P* < 0.001; ns, not significant.)

In addition to punishment US, we also tested the priming effect of reward US by optogenetically activating sugar sensory neurons in flies starved for two days. We selected SS87269, a Gr64f-split-GAL4 line that labeled labial and tarsal sugar sensory neurons but not off-target Gr64f neurons, to express UAS-CsChrimson due to its induction of more robust appetitive memory in flies compared to Gr64f-GAL4 (Shuai et al., 2025). With the same priming and mock training paradigm, we found that flies with their sugar sensory neurons activated during training showed a lower preference for odor OCT than the mock-trained flies (Figure 2A). The ΔPI, also calculated as primed PI minus mock-trained PI, was a negative value that quantified the priming effect of reward stimuli. It was also significantly different from that of the empty driver controls (Figure 2B). These results further extended our understanding of priming behaviors, which were demonstrated to be bidirectional. Animals after experiencing an US would show altered preference for a neutral stimulus that follows, with the direction of alteration opposite to the valence of the US.

Mock-trained and primed flies were different individuals in the experiments above. To directly measure the change of preference in the same animals, we modified the paradigm so that flies in the arena were tested first for their naïve preference for OCT over pure air (Figure 2C). After that, they rested in dark with air for 6min, which was enough time for the activities of most, if not all, KCs and MBONs to recover to the baseline (Turner et al., 2008; Hige et al., 2015a). Flies then underwent red light training and were tested again for the preference for OCT. Comparing the PI between pre-training and post-training, we found that punished flies (MB320C/CsChrimson) developed stronger preference for OCT, while rewarded flies (SS87269/CsChrimson) showed decreased preference, both consistent with previous observations. Their priming scores of ΔPI, calculated as post-PI minus pre-PI, were both significantly different from that of the control flies which mock-expressed CsChrimson under the empty driver (Figure 2D).

Notably, the priming behavior should not be interpreted simply as conditioning of US with pure air. Flies also showed an altered preference for a novel odor after being trained with a different odor (Figure 1A), though this priming effect with CS+ is weak in the T-maze and not observable in arenas (Figure 2—figure supplement 1A), where longer distances are required for flies to navigate to the most desired quadrants, thus resulting in generally lower PIs than those in the T-maze during conditioning. Taking advantage of the more discrete KC representation patterns of ethyl lactate (EL) and pentyl acetate (PA; Campbell et al., 2013), we trained and tested flies with these two odors instead of MCH and OCT and detected the consistent priming behavior in optogenetically rewarded flies (Figure 2—figure supplement 1B-C). Punishment priming effect was still not detected, probably due to the insensitivity of the optogenetic arena. Taken together, these results confirmed the robust and bidirectional priming behaviors in fruit flies, which altered their preference for a normally neutral stimulus in the direction opposite to the valence of the US that they experienced earlier.

### Reward DANs function during aversive priming

The priming effect is hypothesized to be encoded in the MB and the connecting neurons like olfactory memory. The synaptic plasticity between KCs and MBONs is modulated by the output of DANs. Subsets of DANs are activated by punishment or reward cues, as well as by odors alone (Cohn et al., 2015; Siju et al., 2020). In addition to simply conveying US and CS input, DANs also change their responses to CS+ and CS-after conditioning (Riemensperger et al., 2005; Figure 3—figure supplement 1), and thus are considered plastic when animals learn. Since priming also results in altered odor preference, we hypothesized that priming could affect the odor responses of DANs as well. We performed two-photon live calcium imaging of DANs in individual flies which were tethered to receive odors and leg shocks during the experiments (Figure 3C). We expressed the genetically encoded calcium indicator UAS-GCaMP6s (Chen et al., 2013) in a subset of PAM-DANs labeled by 58E02-GAL4 (Jenett et al., 2012). We chose to image PAM-DANs because they were known to be required for appetitive conditioning (Yamagata et al., 2015) as well as the formation of approach memory to CS-after long-term odor-shock conditioning (Jacob and Waddell, 2020), both of which result in the same valence as shock priming. Specifically, we focused on the axons of DANs which project to the γ-lobe of MB, including the γ3, γ4, and γ5 compartments (Figure 3D). γ-MB is known to encode short-term memory in flies (Blum et al., 2009; Qin et al., 2012; Bouzaiane et al., 2015), which shares the same timescale as priming.

**Figure 3.**
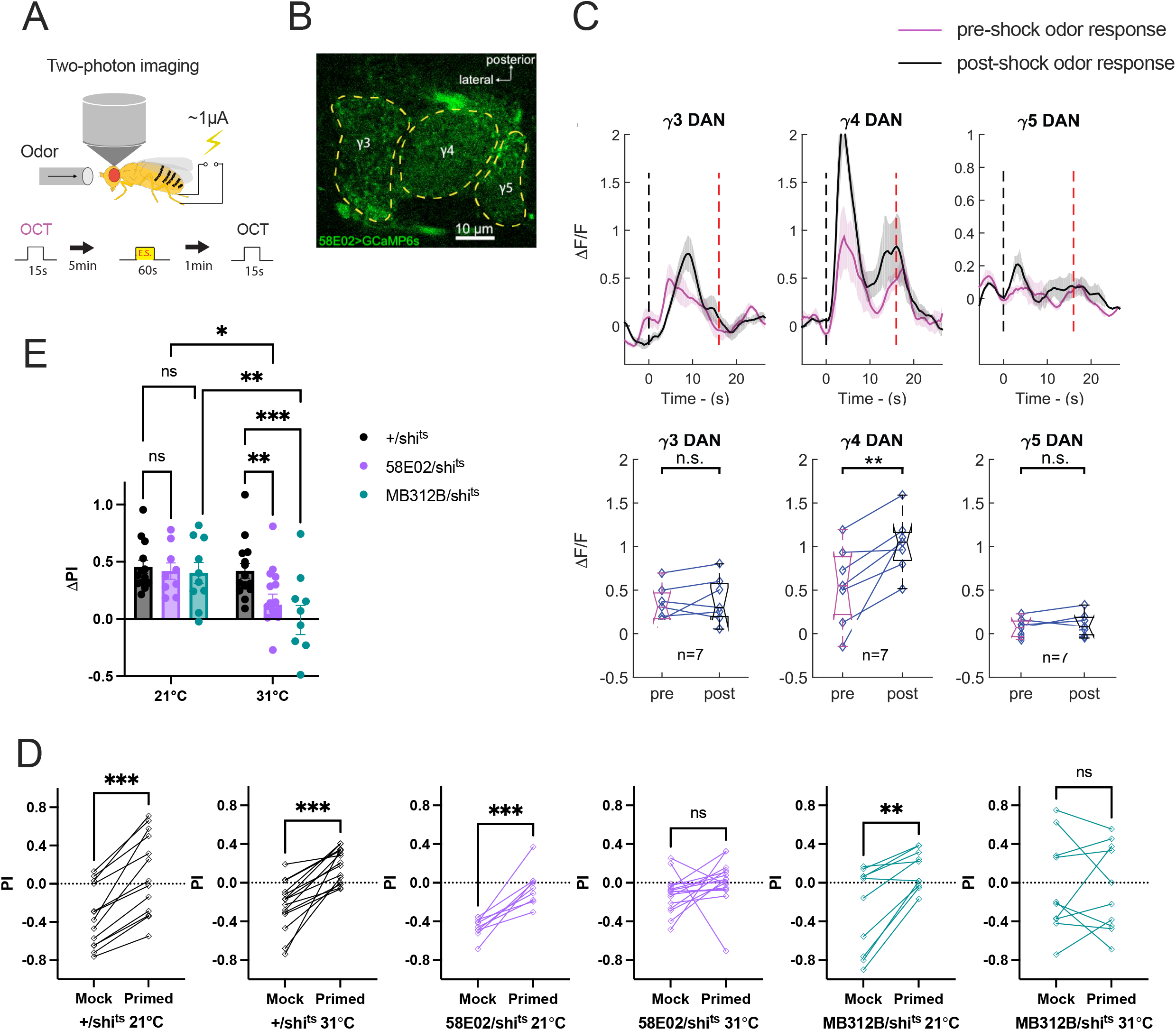
Reward PAM-DANs function during aversive priming. (A) Schematic of the in vivo calcium imaging setup (top) and the experimental protocol (bottom). (B) Sample image of GCaMP6s expressed by PAM-DAN driver 58E02 in γ-MB during odor presentation. (C) γ4-DANs, but not γ3-DANs or γ5-DANs, show increased odor response after shock priming. Top: calcium traces of odor-evoked responses in γ3, γ4, and γ5 compartments before and after electric shocks. Black and red dashed lines denote odor onset and offset, respectively. Data presented as mean (solid curve) ± SEM (shaded area). Bottom: comparison between pre-shock and post-shock odor responses (n=7; paired t-test; ** *P* < 0.01; ns, not significant). (D-E) Silencing reward DANs by *shi*^*ts*^ disrupts aversive priming effect. (D) PI scores of mock-trained flies comparing with shock primed ones, with *shi*^*ts*^ expressed in empty split-GAL4 (+), PAM-DANs (58E02), or in PAM07 and PAM08 (MB312B) at different temperatures (n = 13, 15, 9, 16, 10, 10, respectively; paired t-test). (E) Activation of *shi*^*ts*^ in reward DANs at restrictive temperature disrupts priming (two-way ANOVA). ΔPI = primed PI – mock PI, shown as mean ± SEM (* *P* < 0.05; ** *P* < 0.01; *** *P* < 0.001; ns, not significant).

To compare the calcium activity in DANs responding to an odor before and after priming, we first presented OCT to a naïve fly for 15s to record its pre-shock odor response (Figure 3C). We waited for at least 5min for the odor-evoked calcium activities to fully revert to baseline in the animal, and then shocked the fly for 1min with 12 electric pulses. After another 1min interval, the same odor was delivered to the fly, and the post-shock odor response of DANs was imaged. Comparing pre-shock and post-shock, we found no change in calcium activities in γ3-DANs or γ5-DANs (Figure 3E). However, significantly increased odor response was evoked in γ4-innervating DANs after shock priming. γ4-DANs are known as reward DANs, including PAM-γ4<γ1γ2 (PAM07) and PAM-γ4 (PAM08), both activated by reward cues (Tanaka et al., 2008; Aso et al., 2014a; Cohn et al., 2015; Li et al., 2020). Shock conditioning elevates their response to CS- and reduces that to CS+ (Figure 3—figure supplement 1). Although the odor presented after priming is not considered CS-any more, it is still perceived by flies as an odor with enhanced valence, and therefore the plasticity of these DANs in priming is consistent with that in conditioning.

To test if these imaged PAM-DANs, especially γ4-innervating DANs, are required during punishment priming, we examined the shock priming behavior in the T-maze with temperature-controlled blockage of the synaptic transmission in DANs by temperature-sensitive *shibire* (*shi*^*ts*^; Dubnau et al., 2001; Kitamoto, 2001). We expressed *shi*^*ts*^ in PAM-DANs labeled by 58E02, or specifically in PAM07 and PAM08 together labeled by MB312B (Aso et al., 2014a; Figure 3F-G). At the permissive temperature of 21°C, the experimental flies showed normal priming effects which were similar to that of the control flies (+/*shi*^*ts*^), while at the restrictive temperature of 31°C, both DAN-blocked groups showed significantly reduced levels of priming. In addition, RNAi knockdown of dopamine receptor Dop1R1 in KCs also disrupted the shock priming effects in flies (Figure 3—figure supplement 2; Perkins et al., 2015), confirming the requirement for dopamine release onto KCs in priming. Thus, we concluded that γ4-innervating reward PAM-DANs were necessary for shock priming.

### MBON21 is plastic and required during priming

During conditioning, DANs modulate the plasticity of MBONs to guide the behaviors of flies. Activating attractive MBONs triggers approach behavior in flies, while activating repulsive MBONs causes avoidance (Aso et al., 2014b). In the same MB compartment, pairing an odor with the activation of innervating DANs results in odor-specific synaptic depression between the local KCs and MBONs (Hige et al., 2015a), while backward pairing of DAN activities followed by odor presentation, or strong dopamine release alone in the absence of odor, results in potentiation of the KC-MBON synapses (Cohn et al., 2015; Handler et al., 2019). This time-dependent bidirectional synaptic modulation by DANs underlies the opposite behavioral outcomes of the flies trained by forward and backward conditioning, and thus could also potentially account for the priming behavior. MBONs are classified into 21 types of typical MBONs with dendrites confined to the MB lobes, and 14 atypical MBONs which have their dendritic arbors extending both inside and outside of the MB (Aso et al., 2014b; Li et al., 2020; Rubin and Aso, 2023). To search for the MBONs responsible for priming, we hypothesized that γ4-innervating MBONs could show plasticity during priming like they do in conditioning (Figure 4—figure supplement 1A), considering their direct connections with γ4-DANs. We imaged the dendritic zones of the repulsive MBON-γ4>γ1γ2 (MBON05) before and after electric shocks but found no significant change in their odor responses (Figure 4—figure supplement 2). The adjacent MBON-γ2α’1 (MBON12), MBON-γ3 (MBON08) and MBON-γ3β’1 (MBON09), which were attractive MBONs also known to be plastic during conditioning (Figure 4—figure supplement 1B-C), showed no plasticity during priming either (Figure 4—figure supplement 2).

The only other typical MBON innervating γ4-MB is the understudied MBON-γ4γ5 (MBON21; Li et al., 2020). Activation of MBON21 paired with odor induces aversive memory, and its activation alone drives avoidance in flies (Rubin and Aso, 2023; Mohammad et al., 2024). We imaged this MBON labeled by VT999036 (Figure 4A; Aso et al., 2014b; Shuai et al., 2015; Mohammad et al., 2024) with GCaMP6s pre-shock and post-shock using the same paradigm as DAN imaging, and found that MBON21 in naïve flies didn’t show significant response to aversive chemical odors including OCT or MCH, or to electric shocks alone. However, its calcium level responding to odor presentation was significantly reduced below baseline after shock priming (Figure 4B). Although a negative calcium response to odor is uncommon for MBONs (Hige et al., 2015b), the downregulation found in MBON-γ4γ5 is consistent with the upregulation of γ4-DANs during priming.

**Figure 4.**
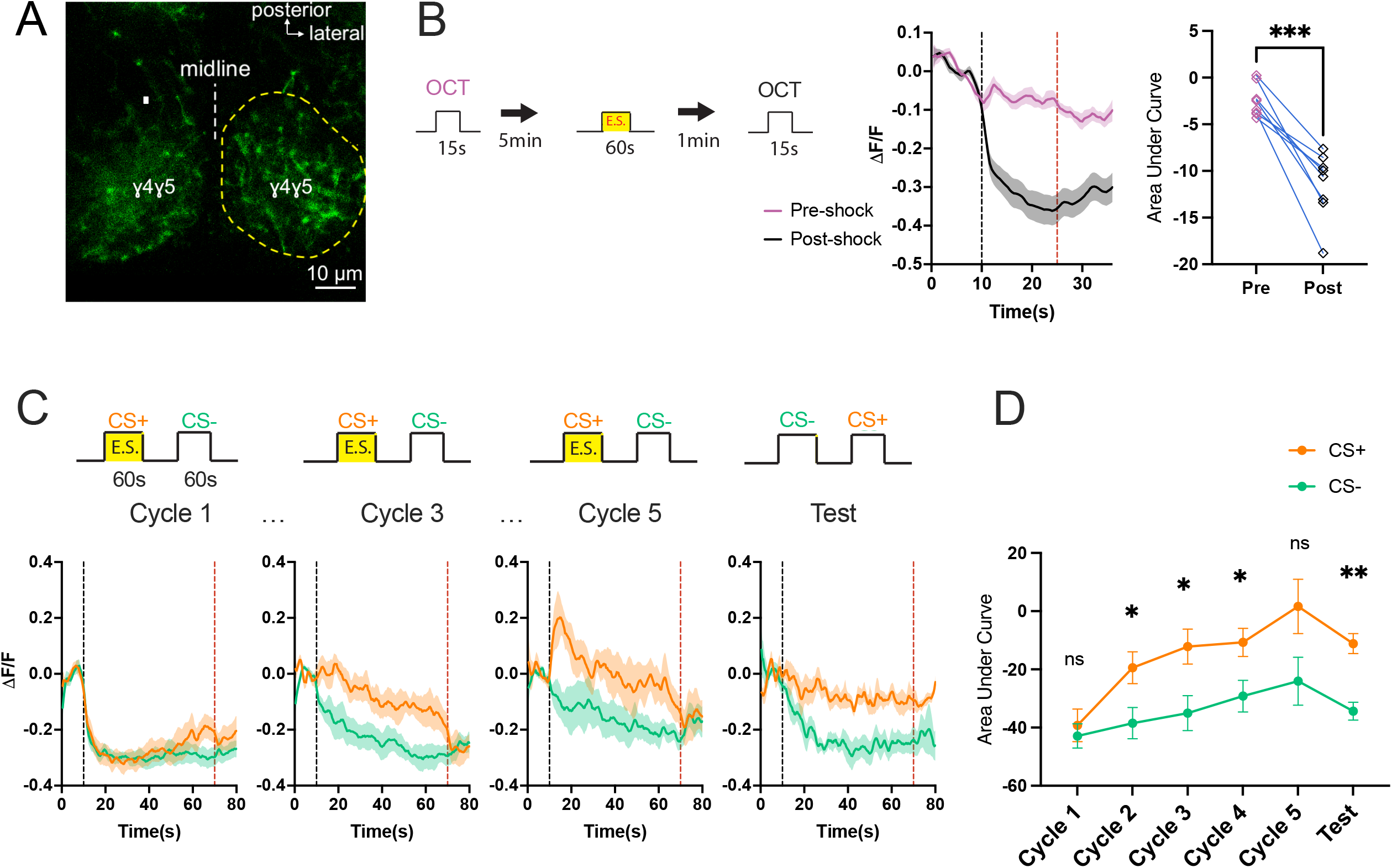
MBON21 shows reduced calcium activity during priming and conditioning. (A) Confocal image of UAS-GFP expressed by MBON21 driver VT999036 in γ4γ5-MB in both hemibrains. (B) Shock priming results in reduced calcium in response to odors in MBON21. Left: schematic of the experimental protocol. Middle: calcium traces of odor response in MBON21 before and after electric shocks. Black and red dashed lines mark odor onset and offset, respectively. Data represented as mean (solid curve) ± SEM (shaded area). Right: comparison between pre-shock and post-shock odor response (n=8; paired t-test; *** *P* < 0.001). (C-D) MBON21 is plastic during spaced shock conditioning. (C) Top: experimental paradigm of 5x shock conditioning and test. Bottom: calcium traces of MBON21 in response to CS+ (orange) and CS-(green) during training cycle 1, 3, 5 and test. Calcium traces of training cycle 2 or 4 are not shown. Black and red dashed lines indicate odor onset and offset time. Data presented as mean (solid curve) ± SEM (shaded area). (D) Comparison between CS+ and CS-responses, shown as mean ± SEM (n=7,6,6,6,5,5, respectively; multiple paired t-test; \**P* < 0.05; ** *P* < 0.01; ns, not significant).

Despite the previous behavioral studies, the physiological change of MBON21 during conditioning was less understood. We imaged MBON21 during 5-cycle spaced shock conditioning, which was a paradigm known to induce long-term memory in flies (Figure 4C). An individual fly was trained and imaged at the same time with shock-odor (CS+) pairing followed by another odor alone (CS-), repeated for five times with long temporal gap in between. Following training, the fly was imaged during a test trial with the two odors without shock. The order of odor delivery during test was reversed to prevent memory extinction resulting from the omission of the expected shock. MBON21 showed decreased calcium levels responding to both odors in the initial cycle. After that, it began to discriminate the two odors and develop a significantly reduced calcium response to CS-compared to CS+ in the later training cycles as well as in the test trial (Figure 4C-D). This plasticity in MBON21 during both conditioning and priming confirms its tuning of repulsion and suggests its necessity in the priming behavior.

We silenced MBON21 by expressing tetanus toxin (TNT; Sweeney et al., 1995) to block its neurotransmission. While the silenced flies still showed detectable priming effect (Figure 5A), their priming score was reduced compared to the controls (Figure 5B), indicating its requirement for normal priming behavior together with other neuronal candidates. To avoid the possible compensation or disruption in the priming circuit resulted from the silenced neurons during early development, we further blocked MBON21 by activating temperature-sensitive *shi*^*ts*^ in adult flies only and observed disrupted priming behavior as well (Figure 5C-D). Because MBON21 functions in priming through down-regulated calcium level according to imaging (Figure 4B), we asked whether artificially activating it by TrpA1, a temperature-sensitive cation channel (Hamada et al., 2008), also resulted in priming defect. Indeed, significantly reduced priming score was found at the restrictive temperature (Figure 5E-F). These behavioral results confirmed the requirement for a normally functioning plastic MBON21 during shock priming. Besides MBON05 and MBON21, there is another MBON that innervates the γ4 lobe of MB, namely MBON29 (MBON-γ4γ5), which is classified as an atypical MBON because it receives synaptic input from outside of the MB as well (Rubin and Aso, 2023). As another avoidance MBON, silencing MBON29 by either TNT (Figure 5—figure supplement 1A-B) or *shi*^*ts*^ (Figure 5—figure supplement 1C-D) also led to reduced priming effect. Taken together, we mapped a priming circuit involving γ4-MB-innervating DANs and MBONs that regulates the olfactory response and memory during the newly observed priming behavior in *Drosophila*.

**Figure 5.**
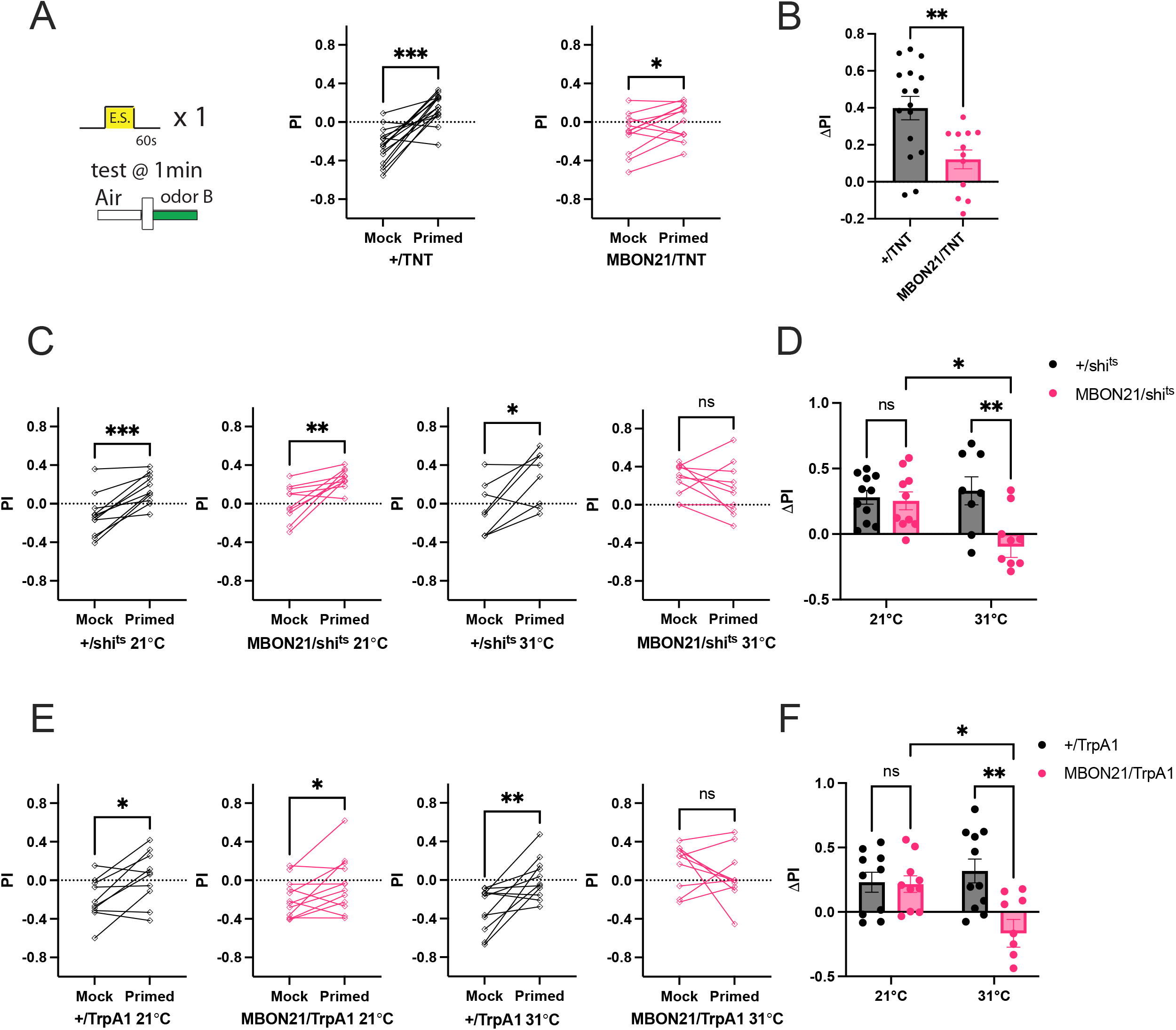
MBON21 is required for priming. (A-B) Silencing MBON21 by TNT disrupts priming effect. (A) The experimental protocol, and the PI scores of mock-trained flies comparing with primed ones (n = 16 and 12; paired t-test). (B) Blocking MBON21 activity by TNT disrupts priming (unpaired t-test). (C-D) Silencing MBON21 by *shi*^*ts*^ impairs priming effect. (C) PI scores of mock-trained flies comparing with primed ones (n = 11, 10, 8, 10, respectively; paired t-test). (D) Activation of *shi*^*ts*^ in MBON21 at restrictive temperature disrupts priming (two-way ANOVA). (E-F) Activating MBON21 by TrpA1 disrupts priming effect. (E) PI scores of mock-trained flies comparing with primed ones (n = 10, 12, 11, 10, respectively; paired t-test). (F) Activation of TrpA1 in MBON21 at restrictive temperature disrupts priming (two-way ANOVA). All ΔPI = primed PI – mock PI, presented as mean ± SEM (* *P* < 0.05; ** *P* < 0.01; *** *P* < 0.001; ns, not significant).

## Discussion

The priming effect characterized in this study has not been observed in flies before. Priming is different from backward conditioning, in which animals form associations between an odor and the relief from punishment perceived as a rewarding cue (Tanimoto et al., 2004; Niewalda et al., 2015; Handler et al., 2019), because the CS used for backward pairing is not present in the priming training protocol. We observed that after being shocked in air, more flies turned towards the normally undesired odor immediately after sensing it in the T-maze compared to unshocked flies. For flies primed in arenas with optogenetic activation (Figure 2C-D), we tracked their real time PI scores during the pre-priming and post-priming odor choice tests, and found that flies after light activation began to show altered PI scores immediately after their initial exploration of the novel odor (Figure 2-figure supplement 2A-B). The calcium responses of γ4-innervating DANs and MBON21 after shock priming also rose to maximum or dropped to the bottom, respectively, right after odor onset with an immediate single peak (Figure 3E and 4B). This suggests that the odor preference change does not depend on a backward conditioning that associates the previous shocks with the odor which only starts to be present at the beginning of the test. In fact, no odor is required to be conditioned during training, but the valence of a novel odor, independent of its identity, is changed by the punishment or reward experience alone. Another behavioral effect, namely blocking odor response by electric shock (BOBE; Song et al., 2018) has been previously reported, where flies staying in a T-maze containing chemical odor and pure air in each arm were shocked before showing enhanced preference for the odor in the following test, and was explained by the interruptive effect of electric shocks on odor responses. While it was distinct from the priming behavior due to the involvement of odor-shock pairing, their results were consistent with and could be partially explained by the priming effect, considering that most flies were shocked in the T-maze arm containing air during BOBE training.

In addition, the priming effect is not due to flies conditioned to associate shocks with pure air. First, sensation of air isn’t known to be encoded by specific neurons in the olfactory sensory system, as flies are raised in air constantly. Even during odor delivery, 99.9% of the diluted odor stream consists of pure air, and thus any conditioning with air would be universally present. Secondly, even if conditioning with air could be contributing partly to the reduced preference for air in the shock priming paradigm without odor (Figure 1B), the paradigm recruiting odor pairing still results in a significant priming effect (Figure 1A). Similarly, appetitive priming effect could also be reliably induced by activating reward sensory neurons together with the presentation of an irrelevant chemical odor (Figure 2—figure supplement 1B-C). Thus, we conclude that the change of odor preference in flies found during priming results from the unconditioned experience instead of another form of conditioning.

In this study, we characterized the functions of γ4-innervating DANs and MBONs which constitute a neural circuit mediating priming effects. Our findings reveal that MBON-γ4γ5 (MBON21) exhibits altered odor response properties following shock stimuli, whereas MBON-γ4>γ1γ2 (MBON05) demonstrates no such plasticity. This differential response pattern suggests different plasticity modulation rules within the same MB compartment and needs further investigation. According to the connectome data acquired by electron microscopy, MBON21 forms multiple feedback circuits onto PAM07 and PAM08 with only one neuronal relay in between, including MBON05, PPL101, APL, and FB4R (Zheng et al., 2018; Li et al., 2020; Hulse et al., 2021). These connections could serve as the potential pathways to complete the feedback loop from MBON-γ4γ5 to γ4-DANs, which show plasticity during priming.

The concept of priming has been used almost exclusively to explain human behavior in the past. Unlike the limited number of studies on priming in other animals that focused on changes in response time or success rates of specific trained behaviors, we found a change in the valence of olfactorily naïve flies as a result of priming without any prior conditioning. This priming behavior, along with its associated neural networks, is possible to be conserved across species, with fruit fly being a simple yet sufficiently complex model organism for investigation.

## Materials and methods

### Fly strains

Flies that were not temperature or light sensitive were reared on standard cornmeal-agar-molasses medium at 25°C and 50% humidity, under 12h light / 12h dark light cycles. Flies expressing shi^ts^ or TrpA1 were crossed and raised at 22°C until experiments in adults. Flies expressing CsChrimson were crossed and raised in dark on standard cornmeal food supplemented with 0.2mM all-trans-retinal (Sigma-Aldrich, St. Louis, MO) until sorted on a cold plate and then kept on 0.4mM all-trans-retinal cornmeal food for at least two days before experiments. All flies were 3-12d old at the time of behavioral or imaging experiments. Fly stocks were all obtained from Bloomington Drosophila Stock Center or Janelia Fly Facility.

### Genotypes used in each figure

Figure 1. 2U, a *w*^*1118*^ (isoCJ1) Canton-S derivative (Tully et al., 1994)

Figure 2. Empty stable split-GAL4 > 20xUAS-IVS-CsChrimson-mVenus in attP18

MB320C > 20xUAS-IVS-CsChrimson-mVenus in attP18

SS87269 > 20xUAS-IVS-CsChrimson-mVenus in attP18

Figure 3. (A-B) 2U > pJFRC100-20xUAS-TTS-Shibire-ts1-p10 in VK00005

OK107 > pJFRC100-20xUAS-TTS-Shibire-ts1-p10 in VK00005

(D-E) 58E02-GAL4 > 20xUAS-IVS-GCaMP6s in attP40

(F-G) 58E02-GAL4 > pJFRC100-20xUAS-TTS-Shibire-ts1-p10 in VK00005

MB312B > pJFRC100-20xUAS-TTS-Shibire-ts1-p10 in VK00005

Empty stable split-GAL4 > 20xUAS-IVS-CsChrimson-mVenus in attP18

Figure 4. (A) VT999036 > UAS-GFP

(B-D) VT999036 > 20xUAS-IVS-GCaMP6s in attP40

Figure 5. (A-B) Empty stable split-GAL4 > UAS-TeTxLC.tnt

SS81353 > UAS-TeTxLC.tnt

(C-D) Empty stable split-GAL4 > pJFRC100-20xUAS-TTS-Shibire-ts1-p10 in VK00005

SS81353 > pJFRC100-20xUAS-TTS-Shibire-ts1-p10 in VK00005

(E-F) Empty stable split-GAL4 > 10xUAS-dTrpA1 in attP16

SS81353 > 10xUAS-dTrpA1 in attP16

Figure 1-figure supplement 1. 2U

Figure 2-figure supplement 1. MB320C > 20xUAS-IVS-CsChrimson-mVenus in attP18

SS87269 > 20xUAS-IVS-CsChrimson-mVenus in attP18

Figure 2-figure supplement 2. (A) MB320C > 20xUAS-IVS-CsChrimson-mVenus in attP18

(B) SS87269 > 20xUAS-IVS-CsChrimson-mVenus in attP18

(C) Empty stable split-GAL4> 20xUAS-IVS-CsChrimson-mVenus in attP18

Figure 3-figure supplement 1. 58E02-GAL4 > 20xUAS-IVS-GCaMP6s in attP40

Figure 3-figure supplement 2. OK107 > 2U

OK107 > BDSC_31765 UAS-Dop1R1-RNAi

OK107 > BDSC_62193 UAS-Dop1R1-RNAi

OK107 > BDSC_51423 UAS-Dop1R2-RNAi

OK107 > BDSC_26018 UAS-Dop1R2-RNAi

OK107 > BDSC_26001 UAS-Dop2R-RNAi

OK107 > BDSC_50621 UAS-Dop2R-RNAi

OK107 > BDSC_31981 UAS-DopEcR-RNAi

Figure 4-figure supplement 1. (A-B) R74B04-GAL4 > 20xUAS-IVS-CsChrimson-mVenus in attP18

(C) R52G04-GAL4 > 20xUAS-IVS-CsChrimson-mVenus in attP18

Figure 4-figure supplement 2. R74B04-GAL4 > 20xUAS-IVS-CsChrimson-mVenus in attP18

R52G04-GAL4 > 20xUAS-IVS-CsChrimson-mVenus in attP18

Figure 5-figure supplement 1. (A-B) Empty stable split-GAL4 > UAS-TeTxLC.tnt

SS77450 > UAS-TeTxLC.tnt

(C-D) Empty stable split-GAL4 > pJFRC100-20xUAS-TTS-Shibire-ts1-p10 in VK00005 SS77450 > pJFRC100-20xUAS-TTS-Shibire-ts1-p10 in VK00005

### Shock priming behavioral experiments with T-maze

Experiments were conducted in a dark room lit only by a dim red light hanging above the T-maze, with 50% humidity and 25°C temperature unless temperature-sensitive flies (21°C as permissive and 31°C as restrictive temperature for TrpA1 or *shi*^*ts*^ crosses) were used. Both males and female adult flies aged 3-10 days were collected and flipped into new food bottles or vials 1-3d before experiments. An automated odor-shock training apparatus and a T-maze (Tully et al., 1994) was built with help from Harvard Center for Brain Science (HCBS) machine shop. For behavioral assays of priming, flies receive 1min of 12x 60V electric shock pulses at legs with or without odor delivered at the same time. 1min (or other interval time as indicated in the figures) after the cessation of shocks, flies were transferred into the T-maze to choose between pure air and a chemical odor (0.1% OCT diluted in paraffin oil was usually used unless specified; Sigma-Aldrich) for 1min. The performance index was calculated as PI = (N1–N2) / (N1+N2), where N1 and N2 are the numbers of flies that stayed in the odor and air arm of the T-maze, respectively, at the end of the test. Priming level was represented by ΔPI, calculated as the difference between the PI of primed and mock primed group of flies which had been raised in the same bottle or vial of food to reduce the bottle effect of the flies’ innate odor preferences.

### Priming experiments in olfactory arenas

Fully automated olfactory arenas were designed and built with the help of jET (Janelia Experimental Technology) team as previously described (Kabra et al., 2013; Aso et al., 2014b). Female adult flies aged 3-10d were sorted on a cold plate and fed on 0.4mM all-trans-retinal food for at least 2d before experiments. Flies with sugar sensory neuron activation and their controls were starved on 1% agar (Sigma-Aldrich) for 2d after 0.4mM retinal feeding and before experiments. In one experiment, approximately 20 freely moving flies were sucked by mild vacuum and put into a circular arena. Flies were trained by pre-designed automatic protocols, usually receiving olfactory and LED stimulation during training, and could walk between quadrants with different odors during tests. Odors were diluted in paraffin oil with the concentrations of 0.1% for OCT, 0.13% for MCH, and 0.01% for PA and EL (Sigma-Aldrich), unless otherwise specified. Air or odors with their volume speed set and stabilized by the mass flow controllers (Alicat Scientific, Tucson, AZ) were simultaneously and continuously delivered into the four quadrants from the corners and removed by vacuum connecting to the center of the arena. The locations of air and odors during tests were swapped throughout the experiments. LED light used for optogenetics in this study was 627nm at approximately 20μW/mm^2^. The flies’ trajectories during the experiments were recorded through a glass lid by a camera (Point Grey, Richmond, Canada) with lens (Kowa, Torrance, CA) and an 820nm longpass filter (Pro Optic, Japan) hanging above at 30 frame/s, lit by 850nm infrared LED backlight from under the arena. Movies were later analyzed with a custom-written ImageJ macro script to calculate the real-time PI scores at every 10 frames, and usually a PI score averaged through the last 30s of test was recorded. Figures were then plotted in GraphPad Prism 10 (GraphPad Software, La Jolla, CA).

### Tethered fly training and calcium imaging

For all *in vivo* calcium imaging experiments, a custom-designed chamber was 3D printed for holding the fly. A 5-10d old adult female fly was anaesthetized on ice and glued to a hole cut out from a small piece of aluminum foil. Wax and bio-compatible Kwik-Sil Adhesive (World Precision Instruments, Sarasota, FL) were used as glue to tether the fly in place. The aluminum foil with fly was then attached onto the imaging chamber before immersed in 0.9x Schneider Insect Medium (Sigma-Aldrich, supplemented with 2mM CaCl_2_ and 4mM NaHCO_3_) to cover the dorsal part of the head and the thorax, while leaving the antennae, the abdomen, and the legs free in the air below the foil. The fly was allowed to fully recover from cold anesthesia before its head capsule was opened carefully by removing the cuticle covering the dorsal head with a scalpel and forceps. Obstructing trachea was removed with fine forceps. The animal’s legs were secured in paraffin wax with tips exposed for wire attachment. Copper wires were placed against one of the legs on each body side, surrounded by 1% agarose gel for conduction of electricity. Kwik-Sil glue was then applied around the agarose gel to prevent it from drying in the air. A stimulator (Grass Technologies S48 Stimulator, Astro-Med, Inc., West Warwick, RI) was used to apply constant electric current of approximately 1μA to the wires to deliver shock pulses to the fly body through a pair of legs.

Odor stimulation was achieved by a custom-built odor mixer with a switch design to ensure maintenance of a constant airflow rate when solenoid valves were switched. A continuous stream (2000mL/min) of humidified air, directed at the fly antennae through a Teflon tubing with 1/8-inch-wide inner diameter, was maintained by mass flow controllers (Alicat Scientific). At a trigger, 10% of air stream was diverted through a vial containing a chemical odorant diluted in paraffin oil (Sigma-Aldrich). Chemical odors of 0.3% OCT and 0.15% MCH diluted in 5mL paraffin oil were used for the imaging experiments.

All calcium imaging experiments were performed using a Zeiss two-photon laser-scanning microscope, LSM 780 NLO (Carl Zeiss Microscopy, LLC, White Plains, NY), with a 20x water-dipping lens. Image time series were scanned in a single focal plane focusing on the MB lobe of one hemisphere of the fly brain with a frequency of 2.5Hz, usually starting from 7s before the odor was turned on and ending at up to 1min after the odor was off.

### Image processing and analysis

Processing of calcium imaging data was done in Fiji / ImageJ (Schindelin et al., 2012). When necessary, to correct for animal movement during live imaging, time series images were aligned using the TurboReg plugin of Fiji. For imaging of DANs and MBONs, regions of interest (ROI) were manually drawn based on clear anatomical segregation of the innervation patterns in different brain compartments.

Fluorescence intensity was then calculated in MATLAB_R2020b (The MathWorks, Inc, Natick, MA) by averaging the fluorescence across the entire ROI at each frame. For plotting calcium traces, ΔF/F in each frame was calculated by subtracting the difference between the fluorescence intensity and the pre-stimulus baseline intensity, which was the mean of 20 frames (about 7s) ending 1 frame before odor onset, then divided by the baseline value. For quantification of the calcium intensity changes in response to stimuli, we either calculated the area under curve over the entire odor presentation session (Figure 4), or averaged the ΔF/F values during the first 8s of odor onset (other imaging plots). Calcium traces and quantification plots were plotted in MATLAB or Prism.

### Statistics

Statistical analysis was performed in MATLAB or Prism. The statistical method, significance level, and sample size used for each experiment were indicated in the corresponding figure legends. Sample size had not been predetermined by pilot experiments.

## Acknowledgements

We would like to thank Edward Soucy (Harvard Center for Brain Science) and Yichun Shuai (Janelia Research Campus, HHMI) for help with the design and manufacture of the T-maze and the behavioral training apparatus, Chuntao Dan (Janelia Research Campus, HHMI) for designing the odor delivery devices, Y. Ding (Harvard) for help with the design of the live imaging chamber, and Yoshinori Aso for providing the behavioral apparatus of the olfactory arena. We thank the Harvard Center for Biological Imaging (RRID:SCR_018673) for infrastructure and support.

## Key resources table

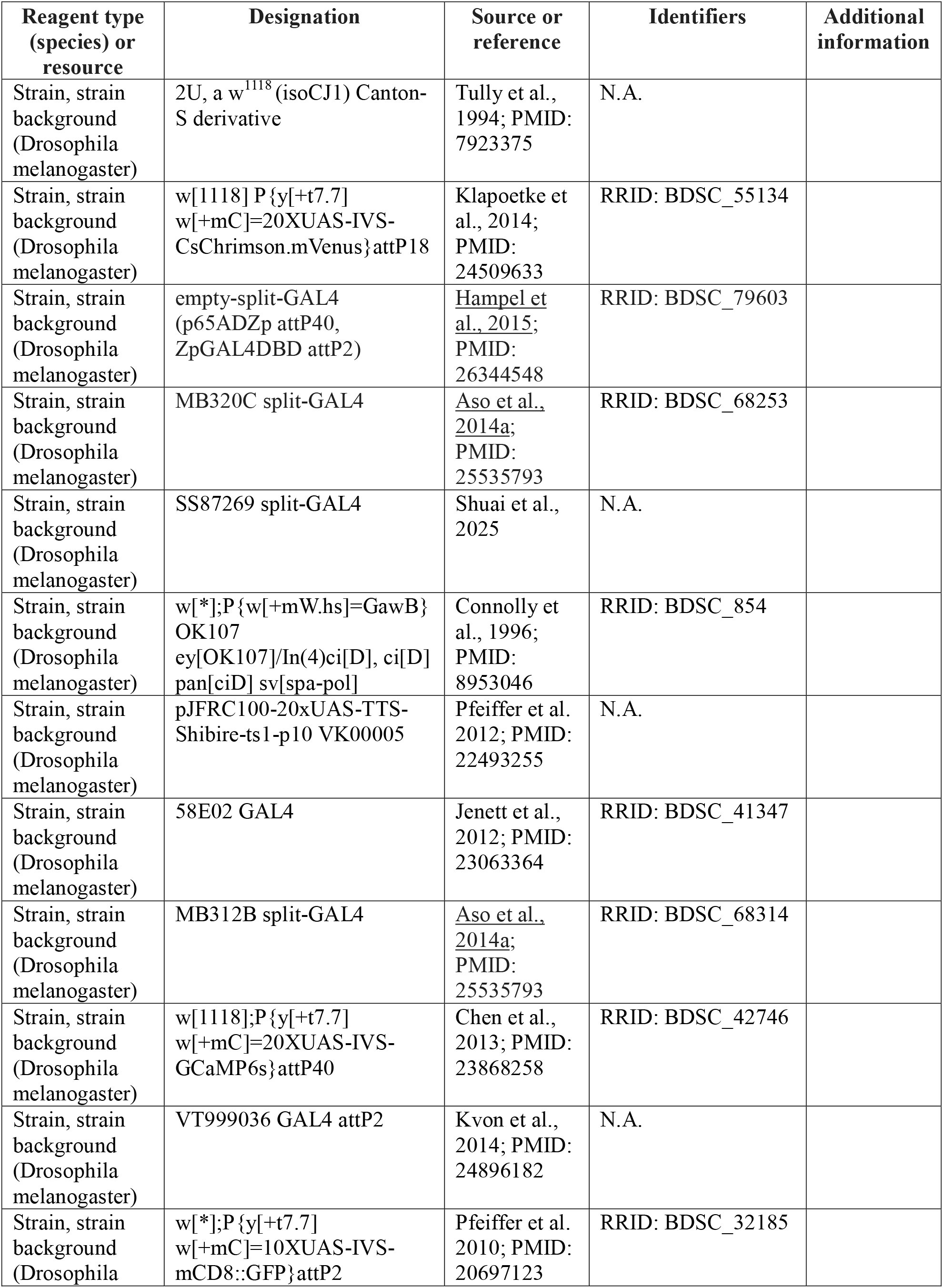
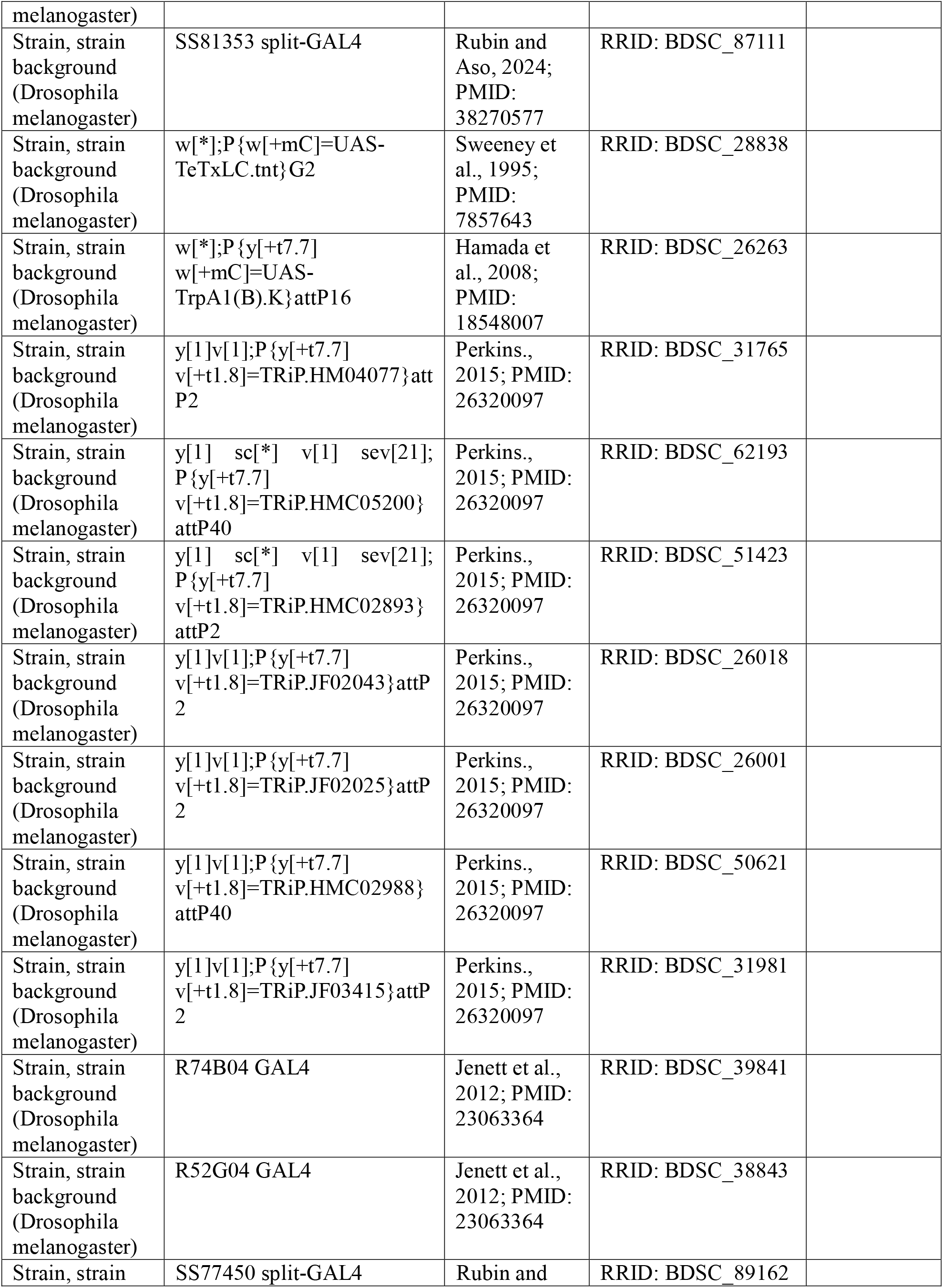
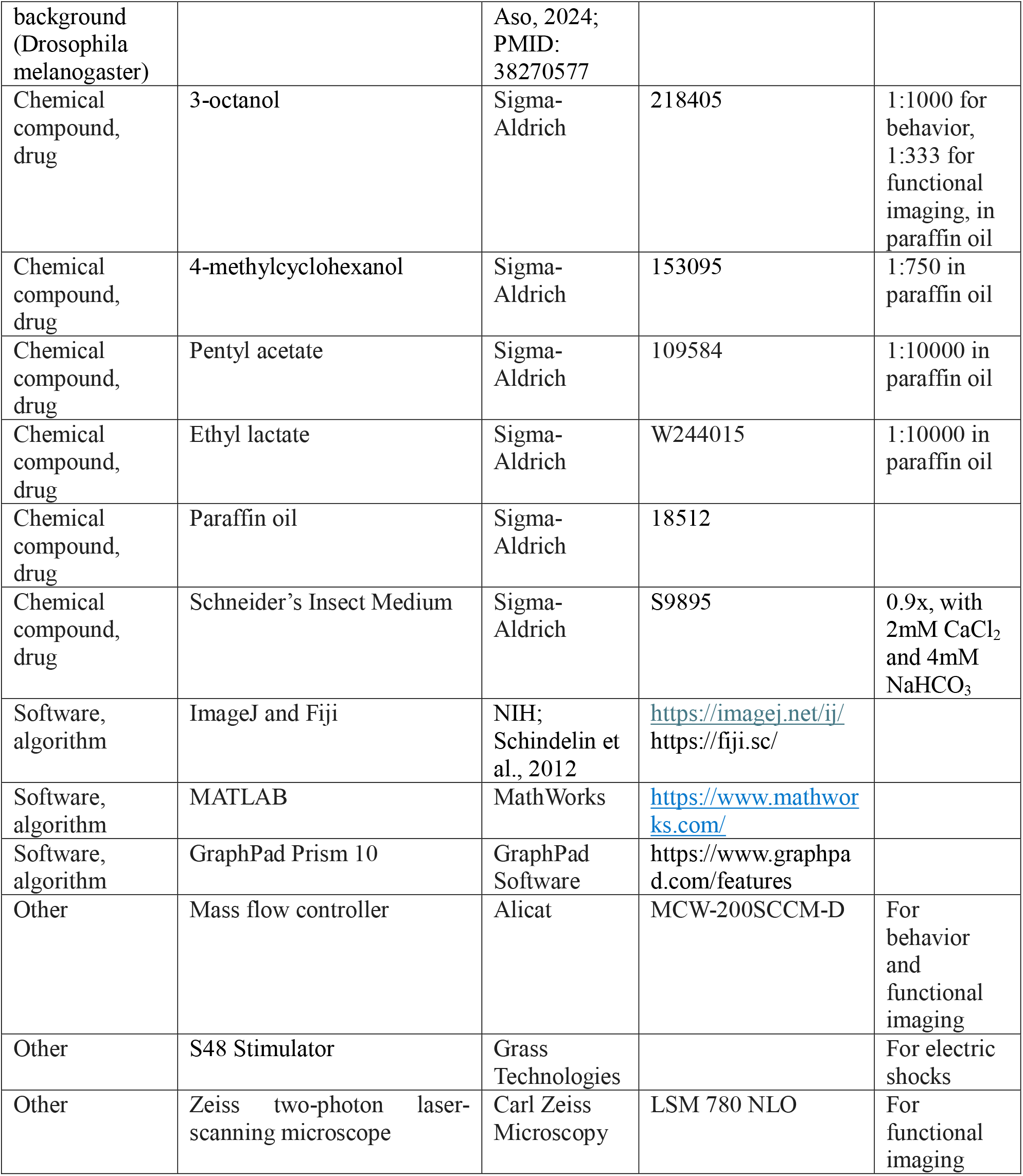

## Figure legends

**Figure 1-figure supplement 1. Aversive olfactory memory is bipartite**.

(A) Schematic of classical spaced aversive olfactory conditioning paradigm. OCT and MCH are used as CS+ in turn in each pair of reciprocal experiments.

(B) Schematic of the paradigm to measure CS-memory component alone. After shock or mock training with CS+, flies are tested for preference between CS- and a novel odor, benzaldehyde (Ben). The CS+ memory component can also be tested in a similar paradigm (not shown), only by switching CS-to CS+ in the test phase.

(C-F) Aversive olfactory training results in attraction to CS-in flies. In each panel, on the top is the diagram of training and test. At the bottom are the corresponding PIs for trained and mock-trained (no shock) flies. (C) Training induces attraction to OCT which is used as CS-compared with Ben as reference.

(D) Training induces attraction to MCH as CS-compared with Ben. (E) Training induces attraction to CS-with pure air as reference. PI scores are averaged over MCH and OCT. (F) Training flies with CS+ in the absence of CS-(OCT) fails to induce attraction to OCT compared to Ben. Sample size n indicated in each panel; paired t-test; * *P* < 0.05; ** *P* < 0.01; *** *P* < 0.001; ns, not significant.

(G-K) Full, attraction (CS-), and repulsion (CS+) memory scores at (G) 10min after 1x shock-odor training, or at (H) 30min, (I) 3h, and (J) 24h after 5x shock-odor training. (K) Time course of the full, attraction, and repulsion memories after 5x spaced aversive olfactory training. For full memory, PI calculated as the absolute value of the usual PI; for repulsion memory to CS+, PI = mock PI-trained PI; for attraction memory to CS-, PI = trained PI-mock PI, so that all the PIs are plotted as positive values, shown as mean ± SEM (all n=8).

**Figure 2-figure supplement 1. Priming with one odor alters flies’ preference for a novel odor under restricted conditions**.

(A) Top: experimental protocol. Odor MCH was present during LED activation. Bottom: PI scores of post-activation tests compared with pre-activation tests (n=24 for MB320C; n=12 for SS87269; paired t-test; ns, not significant). Preference for OCT is not changed by priming with MCH.

(B-C) Priming with odor EL alters flies’ preference for odor PA. (B) Top: experimental protocol, similar to (A) but with odor OCT and MCH substituted by PA and EL. Bottom: PI scores of post-activation tests compared with pre-activation tests in flies with CsChrimson expressed in empty split-GAL4 (+, n=16), PPL1-γ1pedc DANs (MB320C, n=14), and sugar sensory neurons (SS87269, n=16; paired t-test within each genotype).

(C) Activation of sugar sensory neurons but not aversive DANs paired with EL reduces flies’ preference for PA. ΔPI = post PI – pre PI, shown as mean ± SEM (one-way ANOVA; * *P* < 0.05; *** *P* < 0.001; ns, not significant).

**Figure 2-figure supplement 2. Primed flies show altered odor preference starting from odor onset**. PIs in real time of flies reacting to OCT vs. air choice tests pre-priming and post-priming (experimental protocol in Figure 2C). All flies stayed in pure air for 1min until OCT started to fill two diagonal quadrants at Time = 0s (shaded background). After initial exploration of the odor distribution, flies expressing CsChrimson in (A) PPL1-γ1pedc DANs (MB320C, n=6) and in (B) sugar sensory neurons (SS87269, n=6) immediately avoided OCT with PIs dropped at different speeds before and after priming, while (C) control flies (EmptySS, n=6) before and after priming showed similar slopes in PI changes immediately following odor onset.

**Figure 3-figure supplement 1**. γ**3-DANs and** γ**4-DANs show plasticity during conditioning**.

Top: experimental paradigm of 5x spaced odor-shock conditioning and test. Bottom – left: schematic diagrams of the locations of DANs in the MB. Middle: calcium traces of PAM12 and PAM07+PAM08 DANs in response to CS+ (red) and CS-(green) during training cycle 1, 3, 5 and during the test. Black dashed lines indicate odor onset time. Only the traces during the first 30s of odor delivery are plotted, shown as mean (solid curve) ± SEM (shaded areas). Right: quantitative comparison between CS+ and

CS-responses at each cycle. Only the ΔF/F during the first 8s of odor delivery were averaged and plotted, shown as mean ± SEM (all n=11; multiple paired t-test; \**P* < 0.05; ** *P* < 0.01; no stars indicate not significant).

**Figure 3-figure supplement 2. Priming effect with RNAi knockdown of different dopamine receptors in KCs**.

Experimental protocols same as in Figure 1B. # of lines indicate BDSC stock numbers. ΔPI = primed PI - mock trained PI. Data are shown as mean ± SEM (n=12,11,6,6,7,9,6,14, respectively; one-way ANOVA with Dunnett correction; * *P* < 0.05; no stars indicate not significant).

**Figure 4-figure supplement 1**. γ**-MBONs show plasticity during conditioning**.

Flies went through 5x spaced odor-shock conditioning, same as in Figure 4C. In each panel, left: schematic diagrams of the locations of MBONs in the MB. Middle: calcium traces during training cycle 1, 3, 5 and during test. Black dashed lines indicate odor onset time. Only the traces during the first 30s of odor delivery are plotted, shown as mean (solid curve) ± SEM (shaded areas). Right: quantitative comparisons between CS+ (red) and CS-(green) responses in (A) MBON05 (n=12), (B) MBON12 (n=11), and (C) MBON08 and MBON09 (n=7). Only the ΔF/F during the first 8s of odor delivery were averaged and plotted, shown as mean ± SEM (multiple paired t-test; \**P* < 0.05; ** *P* < 0.01; no stars indicate not significant).

**Figure 4-figure supplement 2. Electric shocks don’t affect the odor responses in selected** γ**-MBONs**. Top: experimental protocol. Middle: calcium traces of odor-evoked responses in MBON12, MBON08+MBON09, and MBON05 pre-shock (magenta) and post-shock (black), shown as mean (solid curve) ± SEM (shaded area). Black and red dashed lines denote odor onset and offset, respectively.

Bottom: comparison between pre-shock and post-shock odor responses, quantified by ΔF/F averaged over the first 8s of odor delivery (all n=8; paired t-test; ns, not significant).

**Figure 5-figure supplement 1. Atypical MBON29 is required for priming**.

All experimental protocols same as in Figure 5A. (A-B) Silencing MBON29 by TNT disrupts priming effect. (A) PI scores of mock-trained flies comparing with primed ones (n = 16 and 12; paired t-test). (B) Blocking MBON29 activity by TNT disrupts priming (unpaired t-test).

(C-D) Silencing MBON29 by *shi*^*ts*^ disrupts priming effect. (C) PI scores of mock-trained flies comparing with primed ones (n = 11, 10, 8, 10, respectively; paired t-test). (D) Activation of *shi*^*ts*^ in MBON29 at restrictive temperature disrupts priming (two-way ANOVA). All ΔPI = primed PI – mock PI, presented as mean ± SEM (* *P* < 0.05; ** *P* < 0.01; *** *P* < 0.001; ns, not significant).

