## Supplementary figures and images for "A Priming Circuit Controls the Olfactory Response and Memory in *Drosophila*"

### Figure 1-figure supplement 1

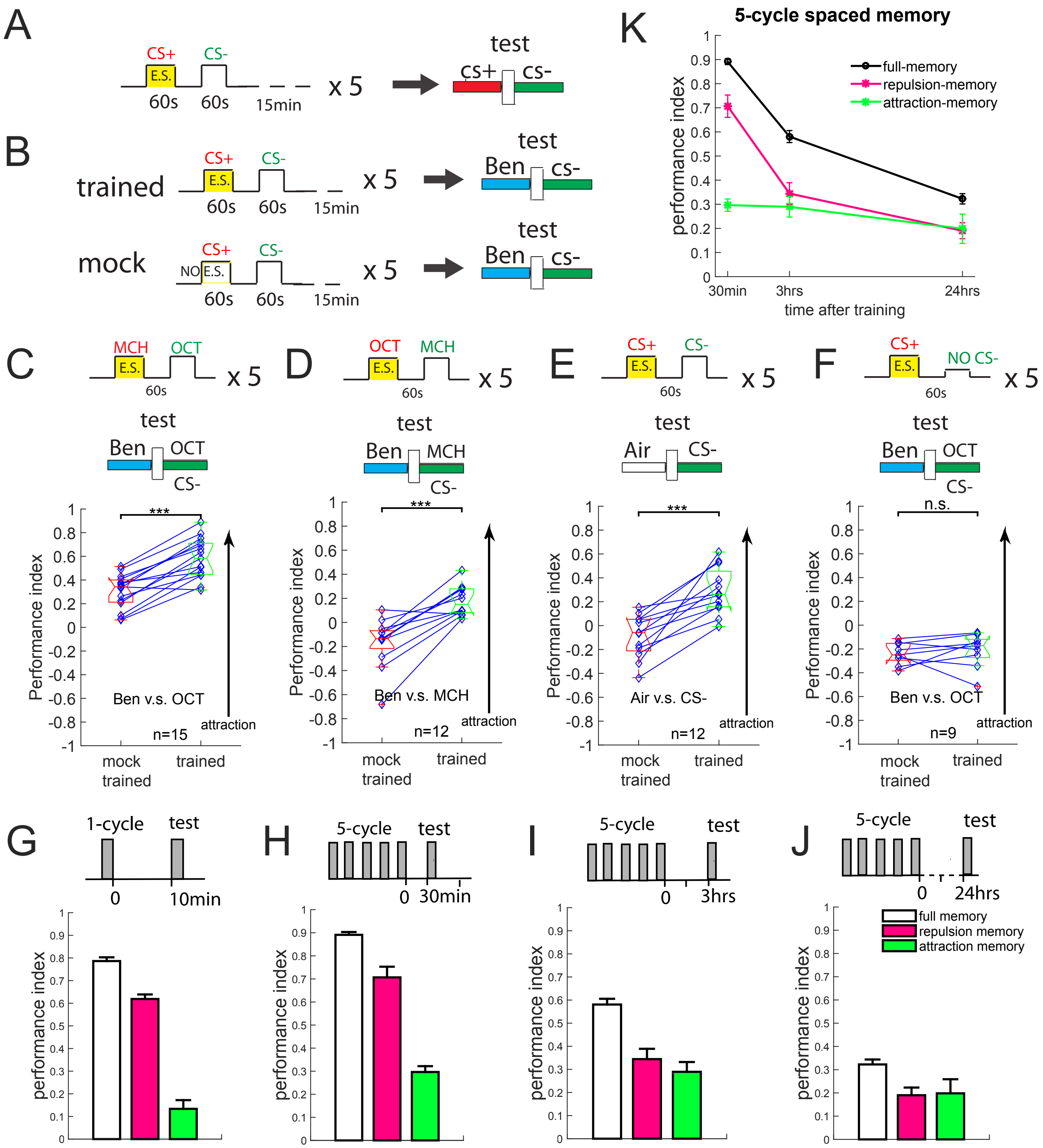

### Figure 2-figure supplement 1

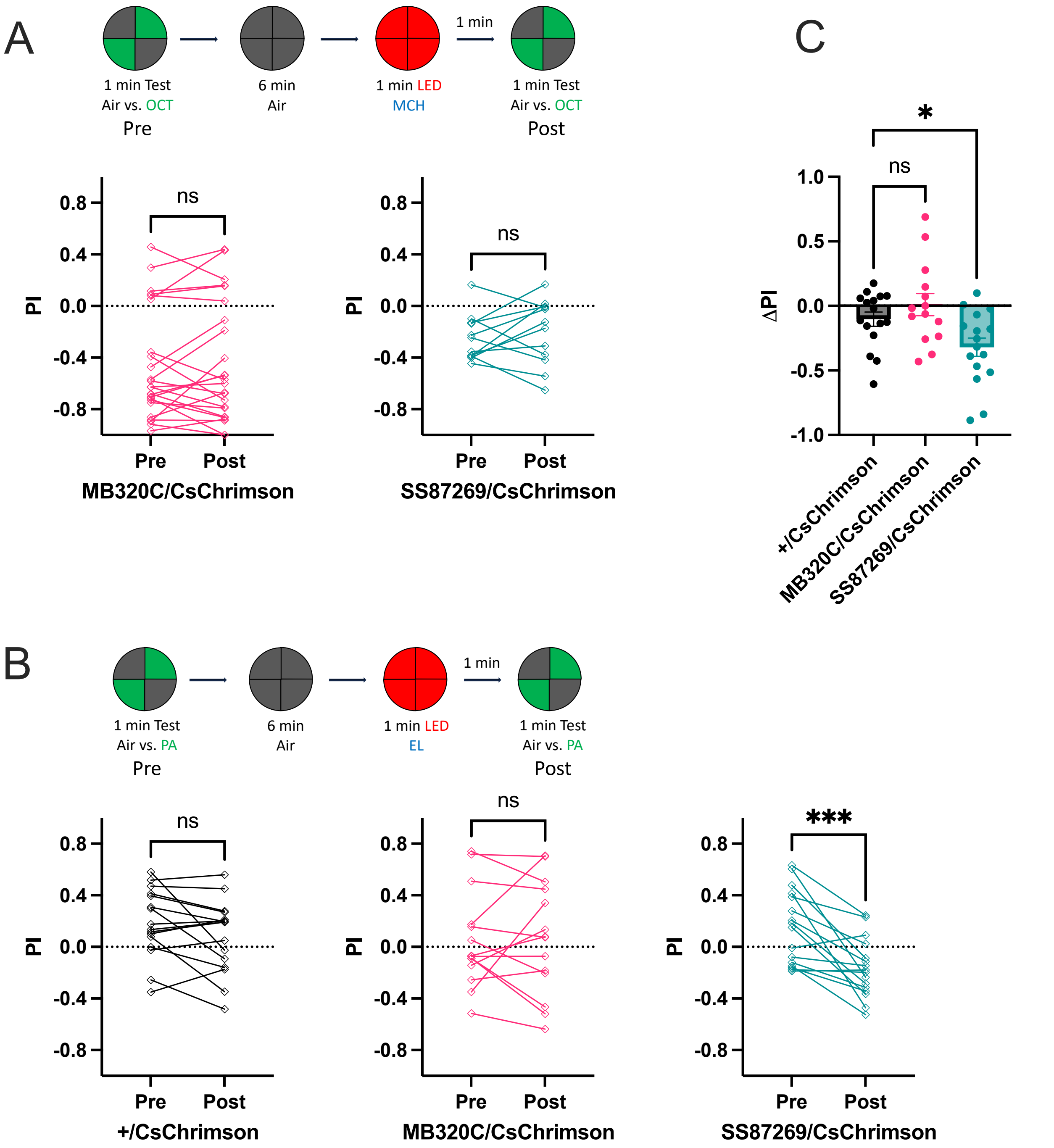

### Figure 2-figure supplement 2

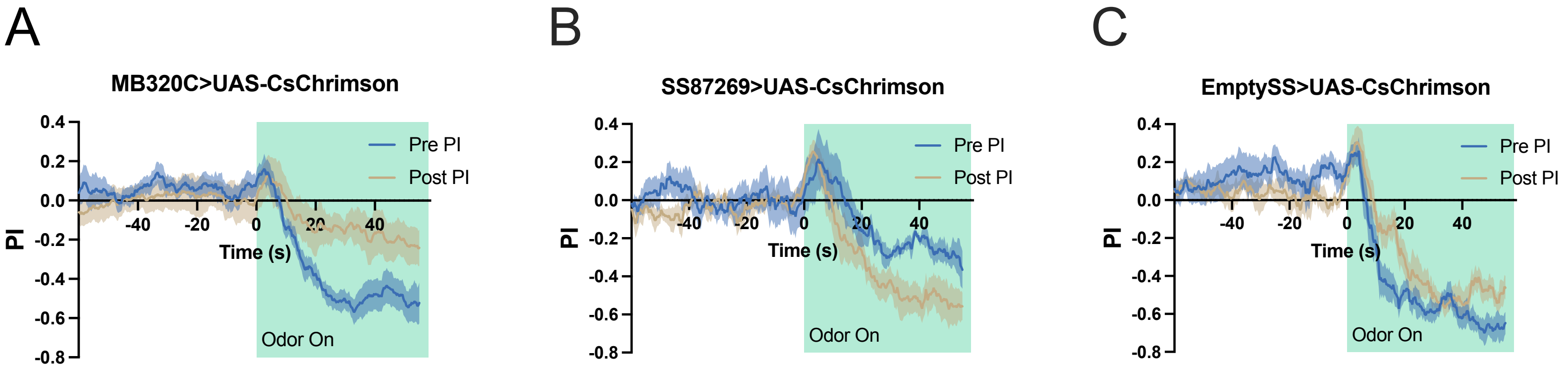

### Figure 3-figure supplement 1

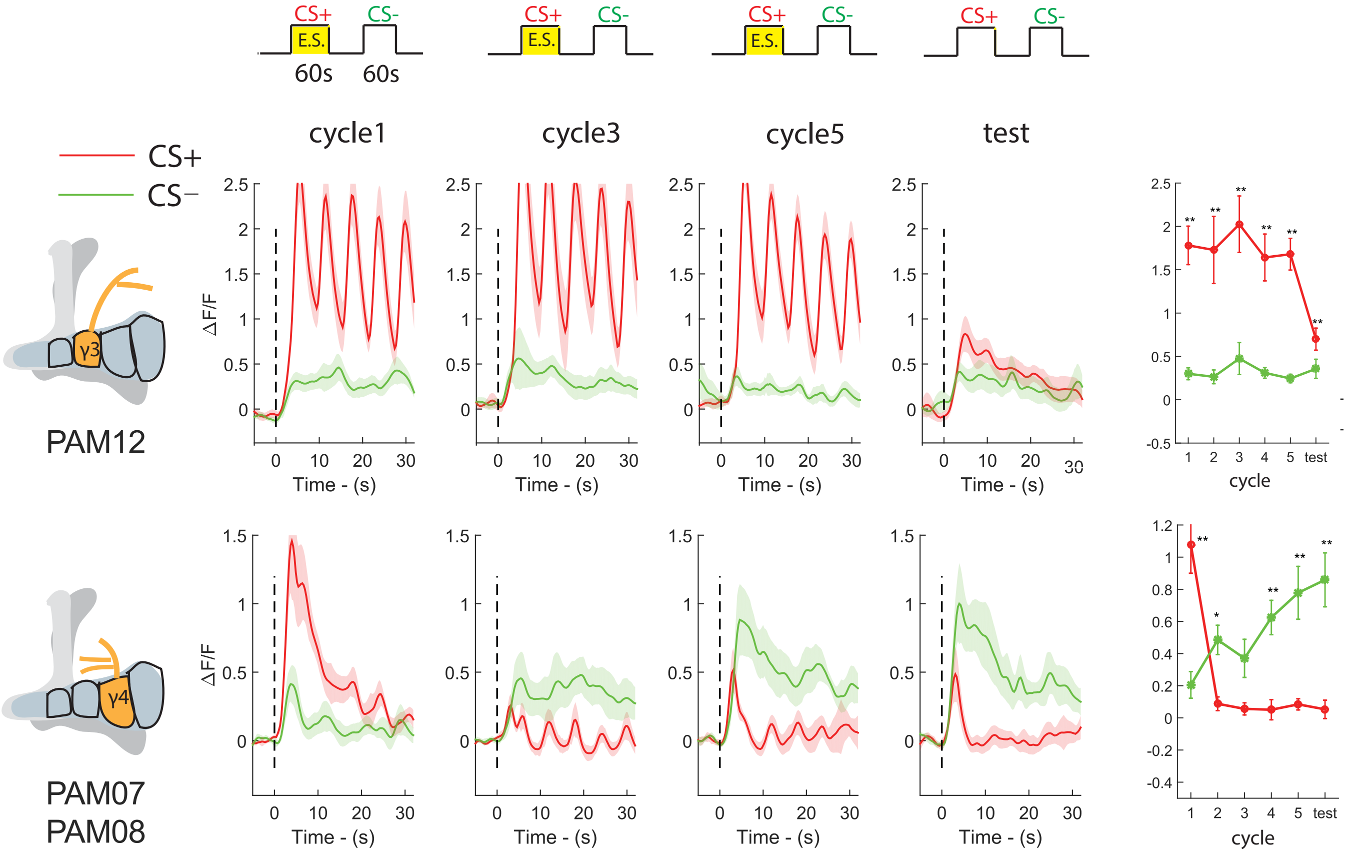

### Figure 3-figure supplement 2

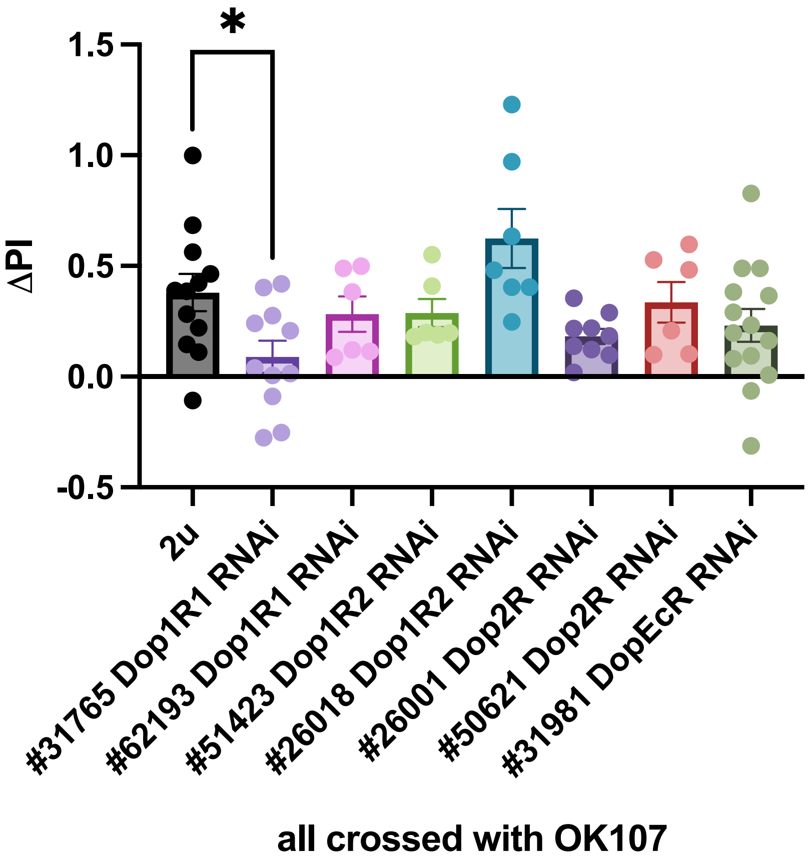

### Figure 4-figure supplement 1

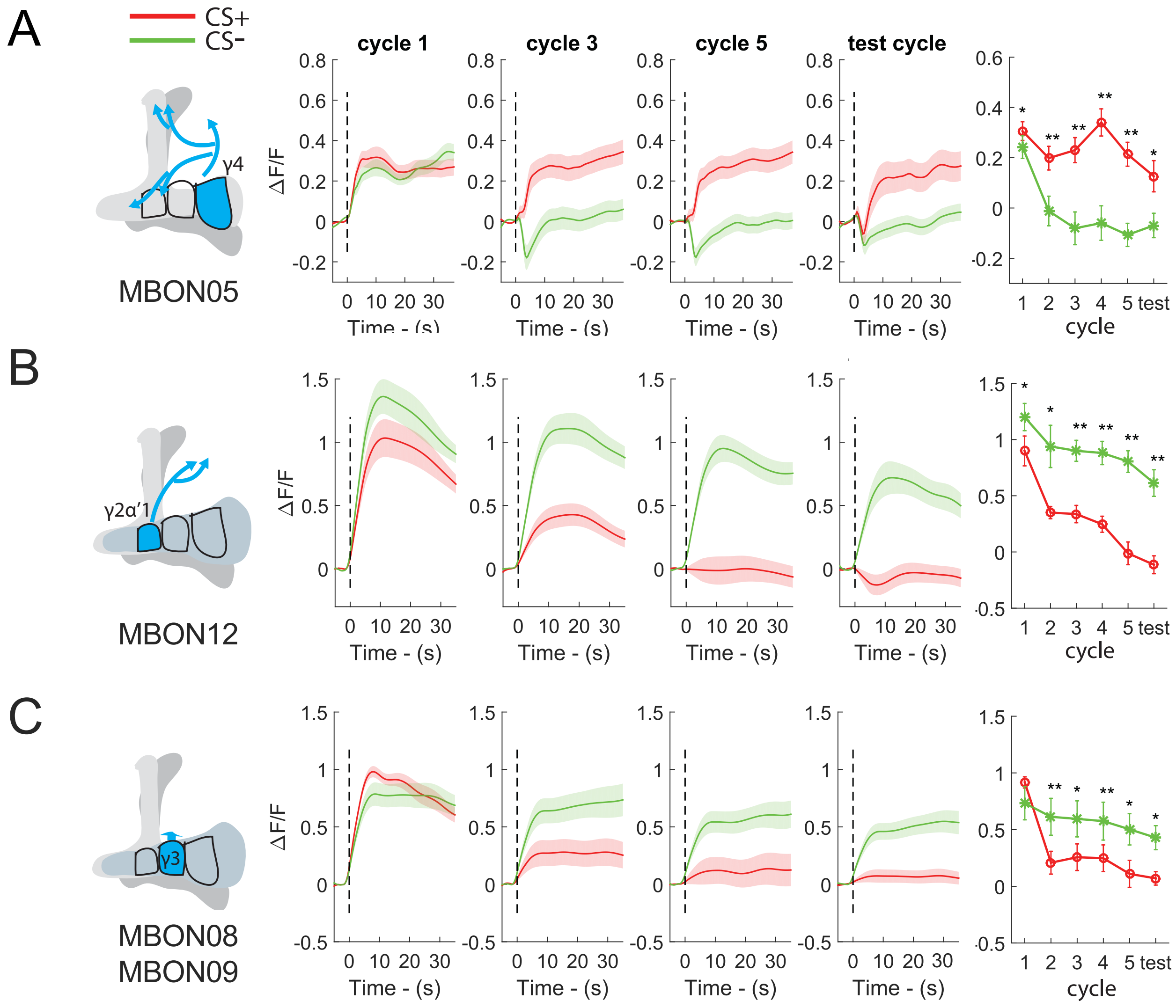

### Figure 4-figure supplement 2

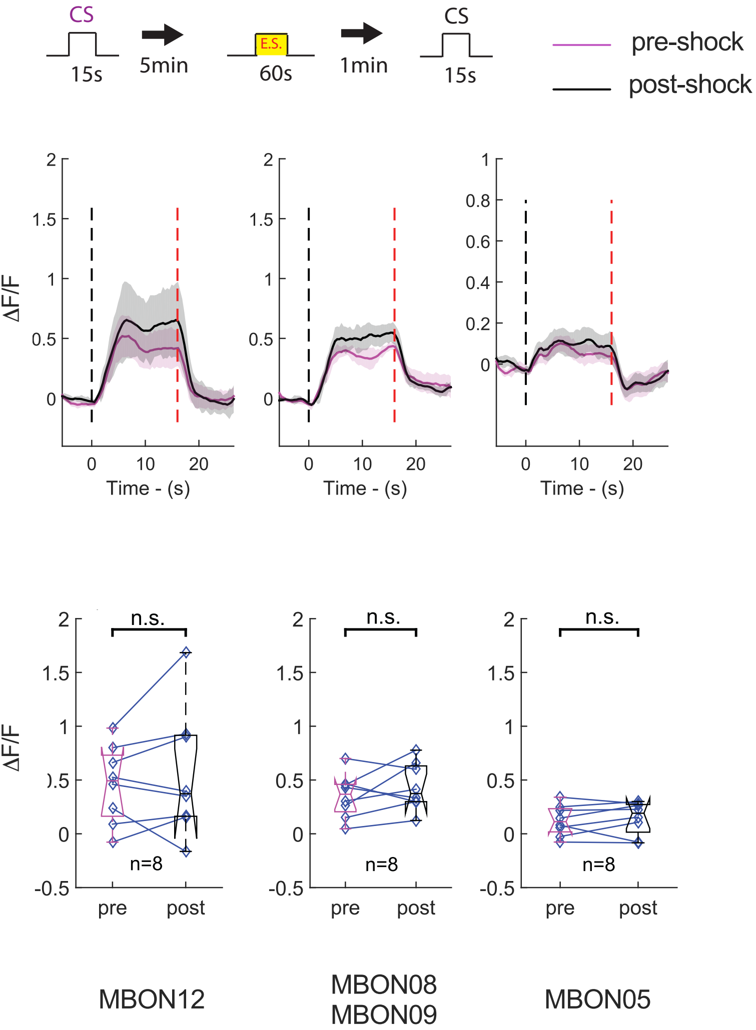

### Figure 5-figure supplement 1

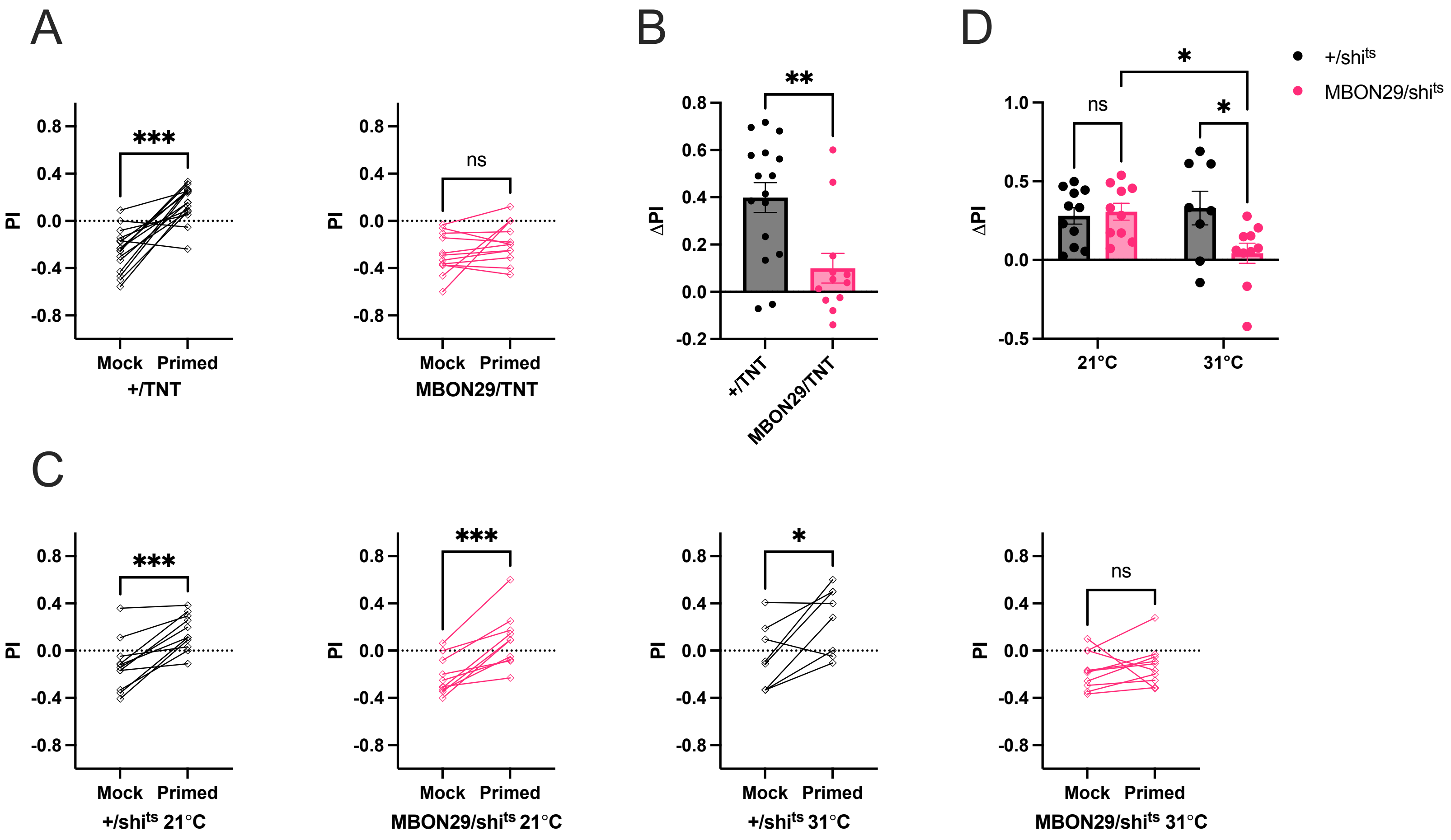
